# Longitudinal stability of the brain functional connectome is associated with episodic memory performance in aging

**DOI:** 10.1101/606178

**Authors:** Olga Therese Ousdal, Tobias Kaufmann, Knut Kolskår, Alexandra Vik, Eike Wehling, Astri J. Lundervold, Arvid Lundervold, Lars T. Westlye

## Abstract

The brain functional connectome forms a relatively stable and idiosyncratic backbone that can be used for identification or “fingerprinting” of individuals with a high level of accuracy. While previous cross-sectional evidence has demonstrated increased stability and distinctiveness of the brain connectome during the course of childhood and adolescence, less is known regarding the longitudinal stability in middle and old age. Here we collected structural and resting state functional MRI data at two time-points separated by 2-3 years in 75 middle-aged and older adults (age 49-80, SD = ± 6.91 years) which allowed us to assess the long-term stability of the functional connectome. We show that the connectome backbone generally remains stable over a 2-3 year time frame in middle- and old age. Independent of age, cortical volume was associated with the connectome stability of several canonical resting-state networks, suggesting that the connectome backbone relates to the structural integrity of the cortex. Moreover, individual longitudinal stability of subcortical and default mode networks were associated with differences in cross-sectional and longitudinal measures of episodic memory performance, supporting the functional relevance. The findings encourage the use of connectome stability analyses for understanding individual differences in cognitive aging. Furthermore, the observation that age-related changes in episodic memory performance relates to the stability of subcortical and default mode networks, provides new longitudinal evidence for the importance of these networks in maintaining mnemonic processing in old age.

## Introduction

Over the past two decades, numerous imaging studies have established a general template of human brain organization. However, few studies have explored the intra-individual variations that lay on top of this blueprint. Recent work on functional brain networks revealed that whereas a substantial proportion of the temporal correlations in the connectome are modulated by cognitive demands, there is also a backbone that remains relatively stable independent of tasks and context (Finn et al., 2015; Kaufmann, Alnaes, Brandt, et al., 2017). The connectome backbone is highly idiosyncratic, allowing the identification of single individuals much like a brain-based fingerprint (Finn et al., 2015; Kaufmann, Alnaes, Doan, et al., 2017; Miranda-Dominguez et al., 2014). Moreover, the connectome stability is sensitive to individual differences in common symptoms of mental disorders in youth (Kaufmann, Alnaes, Doan, et al., 2017), and the neural networks contributing the most to an individual’s connectome– the frontoparietal and the default mode networks (DMN) – are also associated with individual differences in cognitive abilities (Finn et al., 2015). These findings hold great promise for using brain network approaches to advance our understanding of individual variations in cognition and behavior, including an extension to the study of cognitive aging and neurodegenerative disease.

Alterations in the brain grey and white matter structural and functional connectivity are among the hallmarks of cognitive aging (Ferreira & Busatto, 2013; Fjell, Westlye, et al., 2009; L. T. Westlye et al., 2010). These age-related decrements in brain connectivity are paralleled by decline in numerous cognitive functions, likely related to impaired communication between brain regions necessary for maintaining optimal cognition (Ferreira & Busatto, 2013; Fjell, Sneve, Grydeland, Storsve, & Walhovd, 2017; Fjell et al., 2016). Notably, both cortical and subcortical networks are vulnerable to aging (Ferreira & Busatto, 2013; Sala-Llonch, Bartres-Faz, & Junque, 2015), with some networks showing increased and other decreased resting-state connectivity with increasing age (Buckner, 2004; Mowinckel, Espeseth, & Westlye, 2012). Moreover, the extent and rate of change show strong heterogeneity across networks, with frontoparietal and DMN networks, repeatedly identified as the most discriminative of individuals, being particularly sensitive to the aging process (Sala-Llonch et al., 2015). Together these findings suggest that alterations in the structural and functional connectivity of the brain may be related to how well the individual connectome backbone is preserved. Furthermore, it raises the question whether longitudinal stability of the individual connectome is sensitive to concurrent cognitive changes in aging.

Although aging brings about decline in numerous cognitive faculties, episodic memory is one of the most studied. Age-related declines in episodic memory have been reliably identified in both cross-sectional (Hedden & Gabrieli, 2004; Nyberg, Lovden, Riklund, Lindenberger, & Backman, 2012; Ronnlund, Nyberg, Backman, & Nilsson, 2005) and longitudinal (Lundervold, Wollschlager, & Wehling, 2014; Nyberg, 2017) studies of healthy elderly. Moreover, impaired episodic memory is a core symptom of several neurodegenerative disorders, of which Alzheimer disease is the most studied (Gallagher & Koh, 2011). Such age-related changes in memory have been related to altered structural and functional connectivity in prefrontal and DMN networks (Fjell et al., 2015; Nyberg, 2017; Salami, Pudas, & Nyberg, 2014; Staffaroni et al., 2018; E. T. Westlye, Lundervold, Rootwelt, Lundervold, & Westlye, 2011) but more recently also to specific subcortical systems (i.e. the thalamus, amygdala and basal ganglia) (Fjell et al., 2016; Rieckmann, Johnson, Sperling, Buckner, & Hedden, 2018; Ystad, Eichele, Lundervold, & Lundervold, 2010). This is not surprising given the centrality of subcortical nuclei, which are connected to virtually all parts of the cortex (Sah, Faber, Lopez De Armentia, & Power, 2003; Shepherd, 2013), and that neurotransmitters affecting episodic memory target both cortical and subcortical brain structures (Backman, Nyberg, Lindenberger, Li, & Farde, 2006). Moreover, the hippocampal subsection of the DMN and the basal ganglia are conventionally viewed as parallel learning and memory systems (DeCoteau et al., 2007), which may act competitively or cooperatively depending on the context. Accordingly, age-related changes in resting-state functional connectivity of both systems have been linked to deficits during mnemonic processing (Rieckmann et al., 2018; Staffaroni et al., 2018), and disorders known to predominantly target the striatal system have also been associated with profound memory impairments already in the earliest stages of the disease (Solomon et al., 2007).

In the present study, we investigated the longitudinal stability of the connectome backbone in middle and old age, and how this relates to changes in episodic memory and structural indices of aging. We obtained T1-weighted structural and resting-state functional MRI data from 75 middle-aged and older adults (age 49-80, SD = ± 6.91 years) at two time-points separated by 2-3 years, and assessed the longitudinal stability of the whole-brain functional connectome and a set of subnetworks. We hypothesized that connectome stability would decrease as a function of increasing chronological age as well as structural measures of brain aging. Secondly, supported by studies linking age-related cognitive decline to structural and functional connectivity, we hypothesized that weaker connectome stability within subnetworks important for episodic memory would be associated with a steeper memory decline.

## Materials and Methods

### Participants

Healthy volunteers were invited through advertisement to take part in a longitudinal study on cognitive aging involving extensive neuropsychological testing, MRI and genotyping. Participants were assessed up to three times over a period of 6.5 years, of which resting-state functional MRI (fMRI) data were acquired at session 2 (MRI_1_) and session 3 (MRI_2_). MRI_1_ and MRI_2_ were separated by 2-3 years (mean = 2.54, SD = ± 0.28 years). General exclusion criteria included history of substance abuse, present neurological or psychiatric disorder or other significant medical conditions. The protocol was approved by the Regional Committee for Medical and Health Research Ethics of Southern and Western Norway, and all subjects gave written informed consent before participation.

The present study included 75 participants who underwent fMRI at MRI_1_ and MRI_2_. T1-weighted 3D images were evaluated by an experienced neuroradiologist at inclusion, and the presence of brain tumors, cysts, recent infarctions or gross regional or global signal abnormalities lead to exclusion. No participants were excluded based on the neuroradiological evaluation. Moreover, none of the included participants was diagnosed with dementia or mild cognitive impairment (Mini Mental State Exam (MMSE) < 24) (Mungas, 1991). For further participant characteristics, please see Table 1.

**Table 1.**
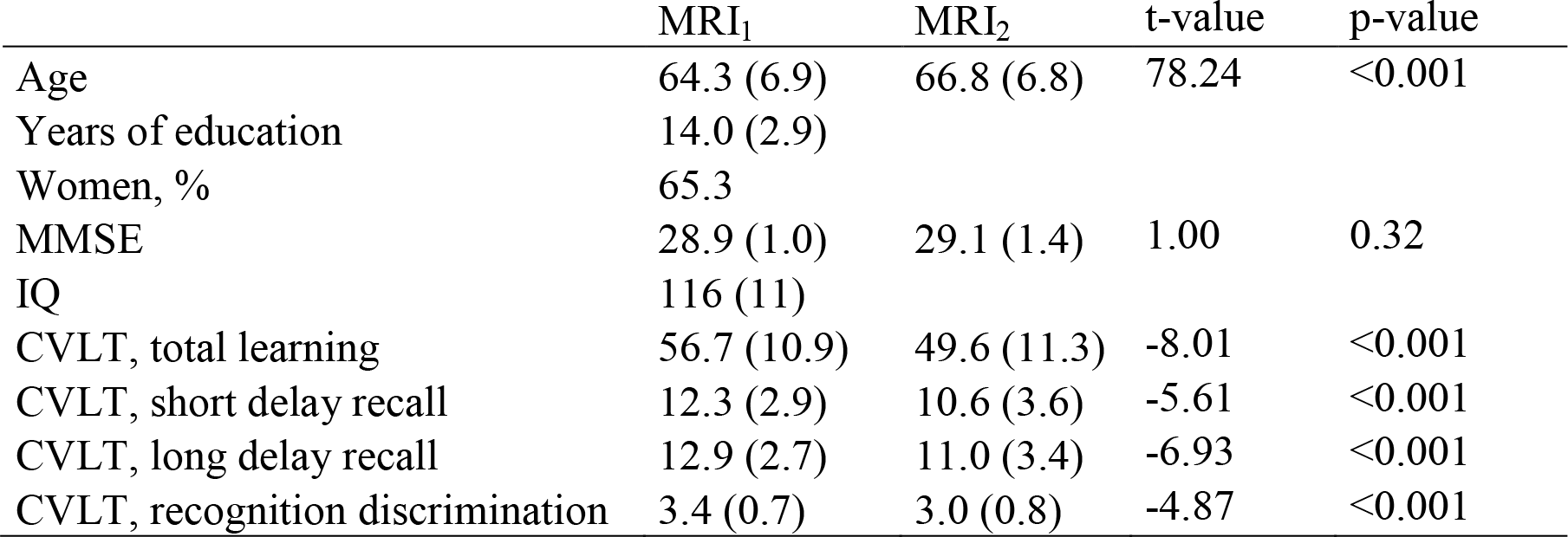
*Means and standard deviation for CVLT-II raw-scores and demographic variables for MRI_1_ and MRI_2_.*

|  | MRI<sub>1</sub> | MRI<sub>2</sub> | t-value | p-value |
| --- | --- | --- | --- | --- |
| Age | 64.3 (6.9) | 66.8 (6.8) | 78.24 | <0.001 |
| Years of education | 14.0 (2.9) |  |  |  |
| Women, % | 65.3 |  |  |  |
| MMSE | 28.9 (1.0) | 29.1 (1.4) | 1.00 | 0.32 |
| IQ | 116 (11) |  |  |  |
| CVLT, total learning | 56.7 (10.9) | 49.6 (11.3) | -8.01 | <0.001 |
| CVLT, short delay recall | 12.3 (2.9) | 10.6 (3.6) | -5.61 | <0.001 |
| CVLT, long delay recall | 12.9 (2.7) | 11.0 (3.4) | -6.93 | <0.001 |
| CVLT, recognition discrimination | 3.4 (0.7) | 3.0 (0.8) | -4.87 | <0.001 |

### Neuropsychological assessments

All participants completed an extensive set of neuropsychological tests at each assessment, and the test scores were evaluated by an experienced neuropsychologist. The battery included tests of executive functions, episodic memory, language, IQ and mental processing speed. Episodic memory function was assessed using the Norwegian translation of the California Verbal Learning Test-Second Version CVLT-II (Delis, Kramer, Kaplan, & Ober, 2000). A list of 16 words (List A) was presented five times, and a total learning score was defined from the sum of correct responses across these trials. Upon completing the fifth trial, a new list was presented (List B), and subjects had to recall the words from list A immediately after List B (short delay free recall). Approximately 20 minutes later, subjects were asked to identify the words from list A again (free long delay recall). Finally, subjects were presented with a larger list that contained items from list A, list B as well as other various distracter items, and asked to identify the 16 items from list A (total recognition discrimination).

The CVLT-II assesses three essential features of episodic memory: learning, recall and recognition, represented by the variables total learning, short and long delay free recall and total recognition discrimination, respectively. Since the episodic memory variables were highly correlated, we used principal component analysis (PCA) to get a compound measure of episodic memory for each subject. In the sample, PCA captured 86.2% of the variance in one single component (PC1). The mean PC1 across MRI_1_ and MRI_2_ as well as the changes in PC1 from MRI_1_ and MRI_2_ were used for the imaging analyses.

### MRI acquisition

Whole-brain, T2*-weighted, echo-planar images (TR= 2000 ms, TE= 50 ms, Flip angle 90º, voxel size, 3.75 × 3.75 × 5.0 mm) were acquired using a GE Signa Echospeed 1.5 T Scanner (General Electric Company; Milwaukee, WI, USA) supplied with a standard eight-channel head coil. A total of 256 volumes (25 axial slices) were acquired, yielding a scan time of approximately eight minutes. Participants were instructed to relax with their eyes closed, to think of nothing in particular, and not to fall asleep. Cushions and headphones were used to reduce subject motion and scanner noise. For anatomical comparison purposes, two T1-weighted 3D inversion recovery-prepared fast spoiled gradient-recalled series (TR=9.11 ms, TE=1.77 ms, Flip angle 7, voxel size 0.94 × 0.94 × 1.40 mmº) were acquired prior to the functional imaging. The imaging parameters were identical for both the T2* and the T1 series at the two time-points. fMRI data from MRI_1_ and MRI_2_ have been previously published (Hodneland, Ystad, Haasz, Munthe-Kaas, & Lundervold, 2012; E. T. Westlye et al., 2011; Ystad et al., 2010; Ystad et al., 2011), however, none of the studies have included longitudinal analyses.

### MRI processing and analysis

T1-weighted 3D MR were processed using the longitudinal pipeline in FreeSurfer v 5.3 (http://surfer.nmr.mgh.harvard.edu), which enables fully automated volumetric segmentation of neuroanatomical structures and longitudinal comparisons. The processing steps included motion correction and averaging, removal of non-brain tissue and automated Talairach transformation. Tessellation of the grey/white matter boundary together with surface deformation following intensity gradients to optimally place the grey/white/CSF borders allowed segmentation of cortex as well as subcortical white matter and deep gray matter structures. All segmented scans were visually inspected.

fMRI data were processed using FMRI Expert Analysis Tool (FEAT), as implemented in FMRIB Software Library (FSL (Smith et al., 2004; Woolrich et al., 2009), (http://surfer.nmr.mgh.harvard.edu)), and included motion correction, spatial smoothing using a six mm full-width at half-maximum (FWHM) Gaussian kernel as well as high-pass temporal filtering (90 s). To minimize the influence of noise (e.g. related to participant motion and vascular artifacts) we applied FMRIB’s independent component analysis (ICA)-based Xnoisifier (FIX (Salimi-Khorshidi et al., 2014)), which uses single-session multivariate exploratory linear optimized decomposition into independent components (MELODIC (Beckmann, DeLuca, Devlin, & Smith, 2005)) to decompose the individual fMRI data sets. Using default options, components were classified as noise and non-noise variability, respectively, using a standard training set supplied with FIX. Components identified as noise and the estimated participant motion parameters were regressed out of the data, and we manually inspected the resulting cleaned fMRI data sets.

The fMRI volumes were registered to the participants’ skull-stripped T1-weighted scans using the FMRIB linear image registration tool (FLIRT, (Jenkinson & Smith, 2001)) implementing boundary-based registration. The T1-weighted volume was nonlinearly warped to the Montreal Neurological Institute MNI-152 template using FMRIB’s nonlinear image registration tool (FNIRT (Anderson, Jenkinson, & Smith, 2007)), and the resulting nonlinear transform was applied to the fMRI data. To control subsequent analyses for data quality and motion confounds, we utilized quality assurance scripts released by Roalf and colleagues (Roalf et al., 2016) and calculated estimates of temporal signal-to-noise ratio (tSNR). One estimate of tSNR per subject and run was calculated by computing voxel-wise mean and SD of the time series (after correcting for linear trends) and averaging the ratio of mean and SD across voxels in the individual brain mask from FSL FEAT. In addition, we estimated an individual mean motion parameter by taking the mean of the relative frame-to-frame displacement (including both rotation and translation) of the raw data.

### Individual level fingerprinting using fMRI data

We used a functional whole-brain atlas consisting of 268 regions of interest (ROIs) (Shen, Tokoglu, Papademetris, & Constable, 2013) and estimated the pairwise Pearson correlations between all ROIs independently for each of the two time points (MRI_1_ and MRI_2_). ROIs were excluded if they were not covered by a minimum of 10% of voxels in all subjects, leading to the exclusion of 20 ROIs in total. The whole-brain connectivity matrix from each individual at each time point was then transformed into a vector of size 1 × 30628 (248 ROIs and 30628 network links between them). Next, we computed the connectome stability in line with the approach by Kaufmann et al. (Kaufmann et al., 2018) which involved computing the within-subject Spearman correlation coefficient between MRI_1_ and MRI_2_ networks.

In addition to parcellating the brain into 248 nodes, we also clustered these nodes based on Yeo et al.’s network scheme (Buckner, Krienen, Castellanos, Diaz, & Yeo, 2011), yielding nine large-scale networks (i.e. medial frontal, frontoparietal, default mode, motor, visual 1, visual 2, visual association, cerebellum, subcortical) (Finn et al., 2015; Kaufmann, Alnaes, Doan, et al., 2017). In line with the whole-brain analysis, we calculated between time points connectome stability scores for each of these nine networks.

### Statistical analysis

All statistical analyses were performed in R (version 3.5.0; R Development Core Team, 2018). Longitudinal analyses modeling the relationship between episodic memory performance and age-, sex, session and time between sessions were performed using linear mixed effects models (lme4 package in R (Bates, Maechler, Bolker, & Walker, 2015)). Separate models were run for each CVLT-II variable (i.e. learning, short delay memory, long delay memory and recognition discrimination). As fixed effects, we entered age (mean across MR_1_ and MR_2_), sex, session and time between sessions (without interaction terms) into the model. In addition to these fixed-effects, the model included subject ID as random factor, modeling the individual level intercept. P-values were obtained by likelihood ratio χ^2^-tests of the full model with the effect in question compared to a model without the effect in question.

Next, we used a general linear model to test for associations between individual whole-brain connectome stability and age while controlling for sex, tSNR, mean motion and time between sessions. The analysis was repeated for each of the nine subnetworks separately, based on studies reporting anatomical differences in the rate and degree of aging (Buckner, 2004). Beyond chronological age, we also tested for associations between cortical or hippocampus volume and connectome stability. As such, the general linear models were expanded to also include a predictor for mean (across MRI_1_ and MRI_2_) or longitudinal changes in total cortical or hippocampus volume while additionally controlling for total intracranial volume (ICV).

To explore the cognitive significance of the whole-brain as well as the subnetworks temporal stability, we used the PC1 obtained from the episodic memory compounds. In separate general linear models we tested for associations between the mean PC1 or changes (across time) in the PC1 and connectome stability, while covarying for sex, age, tSNR, mean motion and time between sessions for each of the nine subnetworks.

To rule out confounding effects of potential extreme values on our results, we excluded subjects with values > 4SD from the group mean from the mixed effects and the general linear models. Throughout the manuscript, we report uncorrected p-values, with a significance threshold for all tests determined by the Benjamini-Hochberg false-discovery rate procedure at q=0.05. In the figures, the regression lines represent the association between dependent and independent variables estimated without covariates.

## Results

### Verbal episodic memory function

Table 1 summarizes the changes in mean scores for all episodic memory measures, supporting significantly lower performance scores in MRI_2_. The contribution of age, sex and session intervals in predicting individual longitudinal episodic memory performance were assessed in linear mixed effect models, with separate models for each of the four CVLT-II variables. Standard likelihood-ratio χ^2^-tests revealed that sex and session were significant predictors of all four memory components (Table 2). As such, performance dropped from MRI_1_ to MRI _2_, and more so for males than females. In addition, higher age was associated with greater decrements of learning, long delay free recall and recognition discrimination (Figure 1).

**Table 2.**
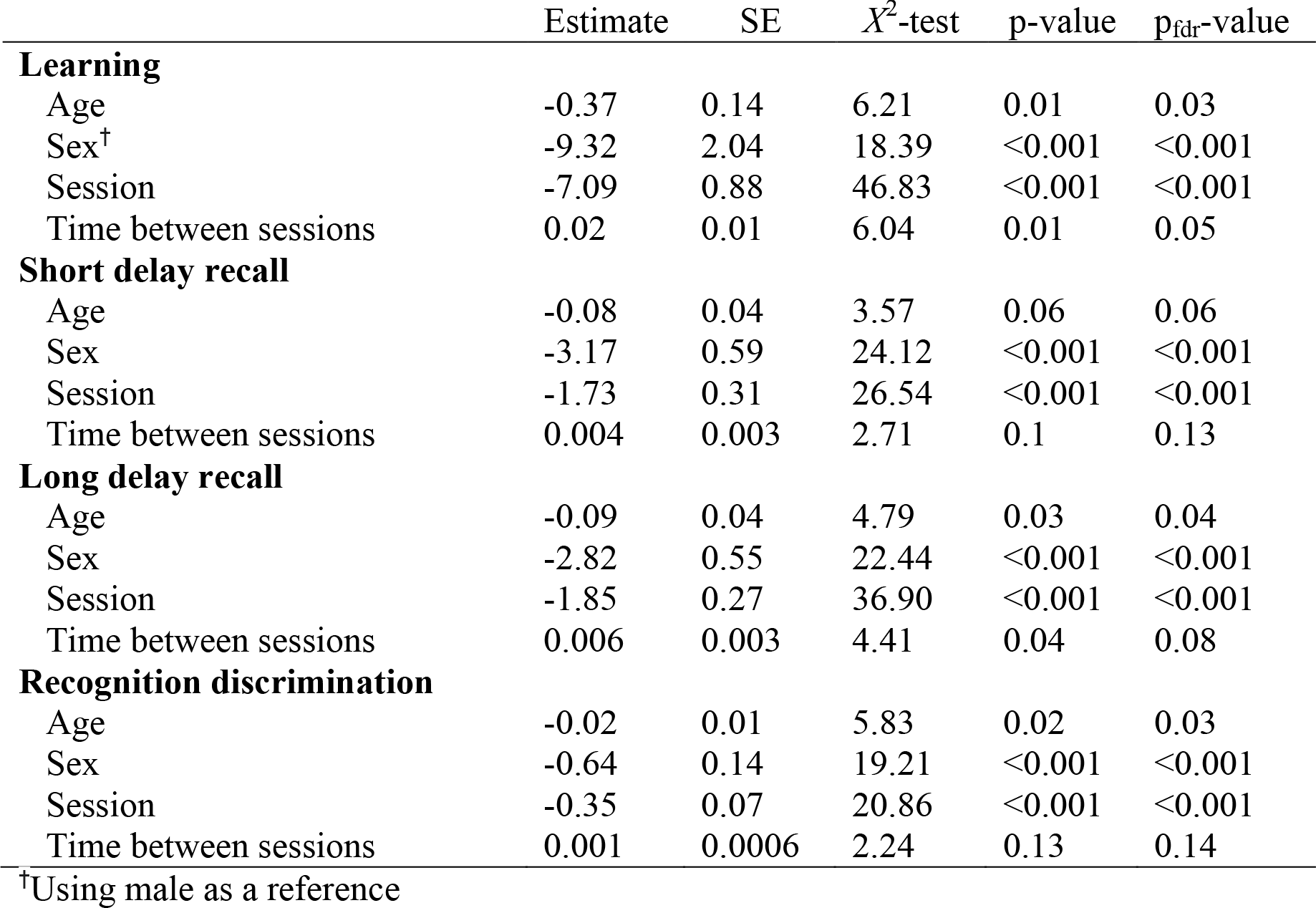
*Association between longitudinal changes in the four different episodic memory components and sample characteristics*.

| | Estimate | SE | $\chi^2$-test | p-value | p<sub>fdr</sub>-value |
| --- | --- | --- | --- | --- | --- |
| **Learning** |  |  |  |  |  |
| Age | -0.37 | 0.14 | 6.21 | 0.01 | 0.03 |
| Sex<sup>†</sup> | -9.32 | 2.04 | 18.39 | <0.001 | <0.001 |
| Session | -7.09 | 0.88 | 46.83 | <0.001 | <0.001 |
| Time between sessions | 0.02 | 0.01 | 6.04 | 0.01 | 0.05 |
| **Short delay recall** |  |  |  |  |  |
| Age | -0.08 | 0.04 | 3.57 | 0.06 | 0.06 |
| Sex | -3.17 | 0.59 | 24.12 | <0.001 | <0.001 |
| Session | -1.73 | 0.31 | 26.54 | <0.001 | <0.001 |
| Time between sessions | 0.004 | 0.003 | 2.71 | 0.1 | 0.13 |
| **Long delay recall** |  |  |  |  |  |
| Age | -0.09 | 0.04 | 4.79 | 0.03 | 0.04 |
| Sex | -2.82 | 0.55 | 22.44 | <0.001 | <0.001 |
| Session | -1.85 | 0.27 | 36.90 | <0.001 | <0.001 |
| Time between sessions | 0.006 | 0.003 | 4.41 | 0.04 | 0.08 |
| **Recognition discrimination** |  |  |  |  |  |
| Age | -0.02 | 0.01 | 5.83 | 0.02 | 0.03 |
| Sex | -0.64 | 0.14 | 19.21 | <0.001 | <0.001 |
| Session | -0.35 | 0.07 | 20.86 | <0.001 | <0.001 |
| Time between sessions | 0.001 | 0.0006 | 2.24 | 0.13 | 0.14 |

<sup>†</sup>Using male as a reference

**Figure 1.**
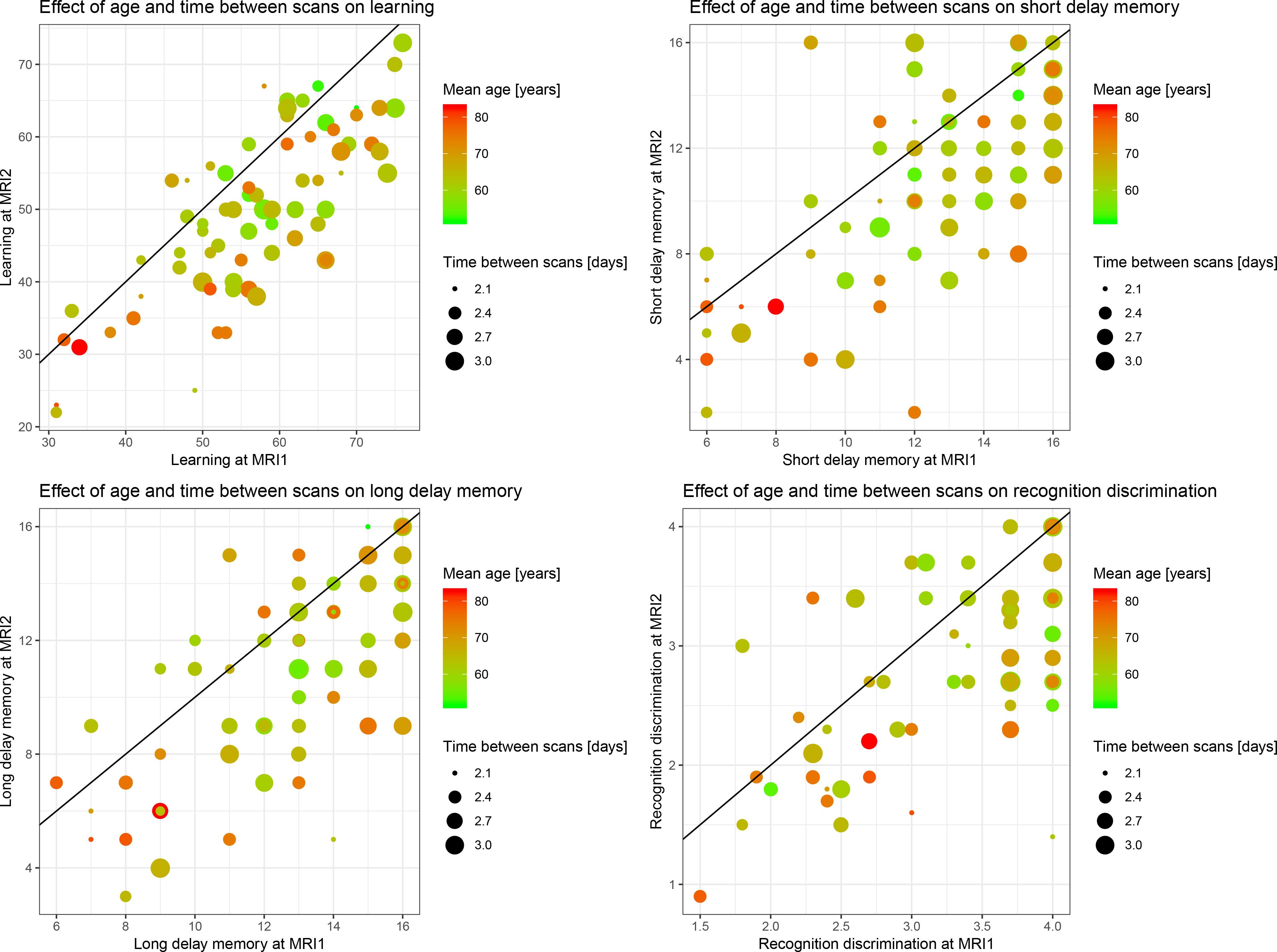
The association between age and time between scans and changes in the four different episodic memory components from MRI_1_ to MRI_2_. The four episodic memory components included total learning, short delay free recall, long delay free recall and total recognition discrimination..

### Functional connectome analyses

Figure 2 shows the results from the connectome stability analyses. In line with recent work in a longitudinal sample of youths (Miranda-Dominguez et al., 2018) the connectome fingerprint retained relatively stable across 2-3 years (mean Spearman correlations between scans for all subjects: rho=0.4, range: 0.15-0.60 for the full brain connectome).

**Figure 2.**
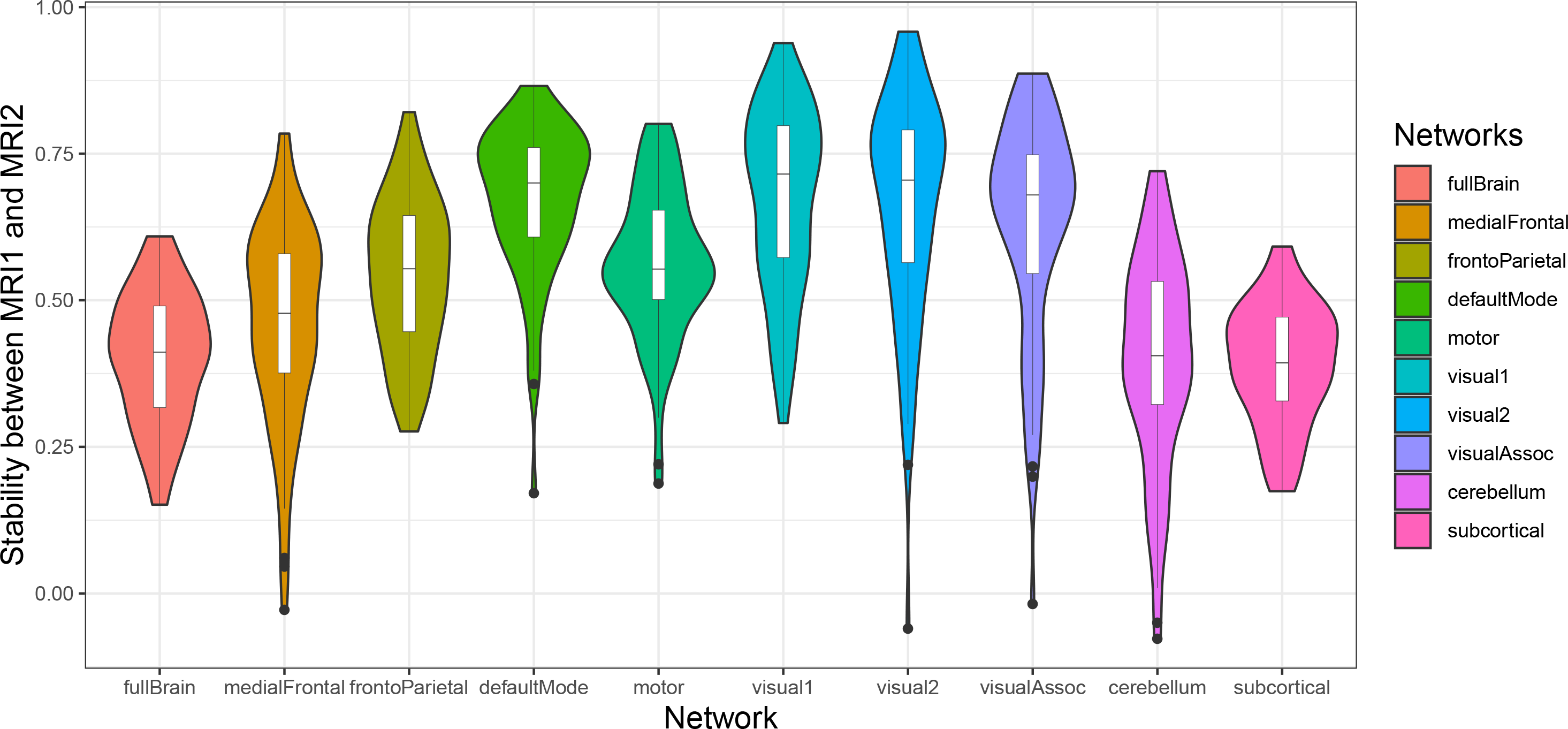
Connectome stability of the whole-brain and the nine subnetworks between the two sessions

Figure 3a illustrates the association between connectome stability and age (mean across MRI_1_ and MRI_2_). The association between age and whole-brain connectome stability (slope (± SE) = −0.003 ± 0.002, t_69_=-1.57, p=0.12) as well as between age and subnetwork connectome stabilities (all p>0.05) were subtle and none remained significant when accounting for multiple comparison.

**Figure 3.**
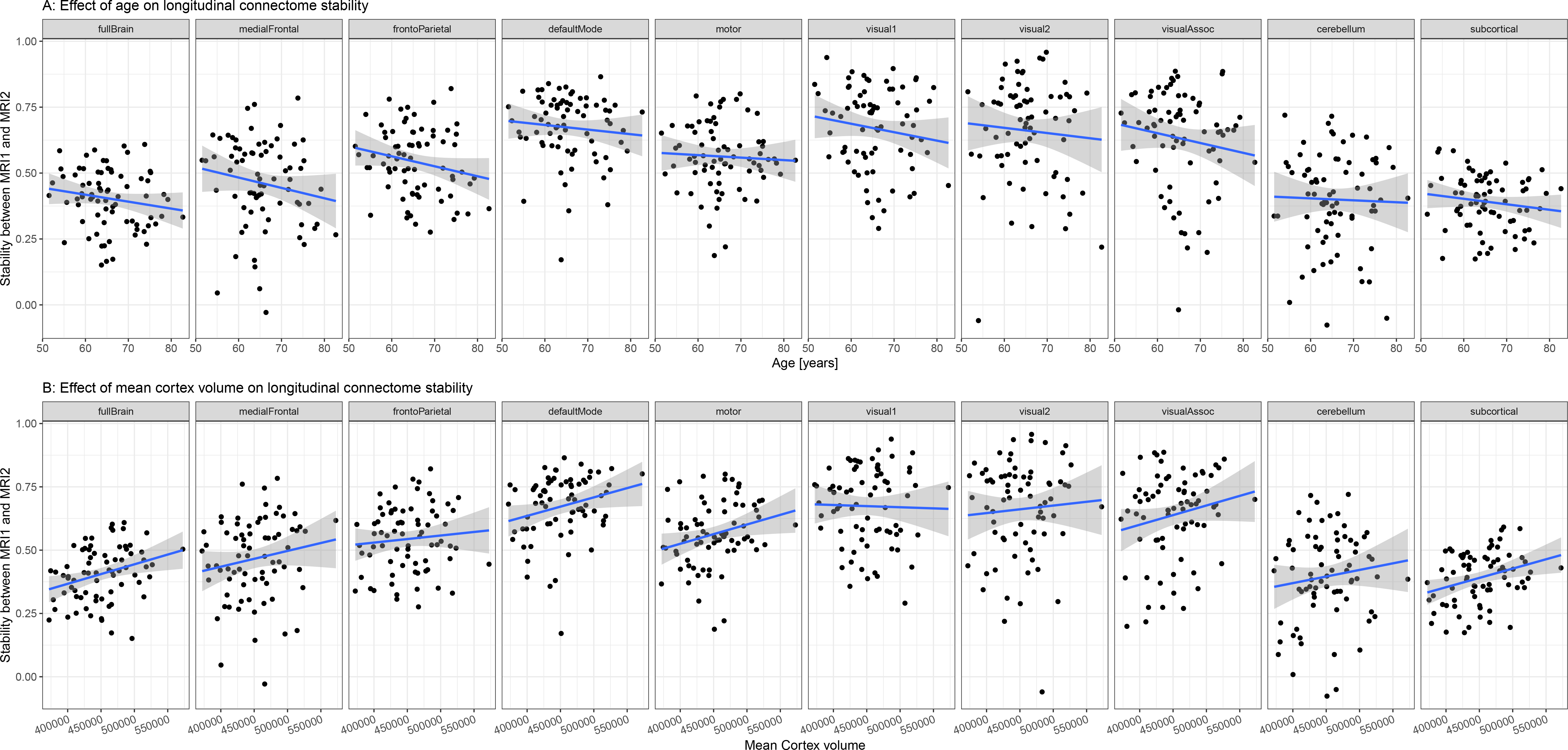
The association between age or total cortical volume and connectome stability. (A) Association between connectome stability and age (mean across MRI_1_ and MRI_2_) for the whole-brain and the nine subnetworks. (B) Association between connectome stability and total cortical volume (mean across MRI_1_ and MRI_2_) for the whole-brain and the nine subnetworks.

We next investigated if the association between connectome stability and age was influenced by cortical volume (defined as CortexVol in FreeSurfer). As such, the general linear models were expanded to also include a predictor for mean (across MRI_1_ and MRI_2_) or time-dependent changes in total cortical volume while additionally controlling for ICV. Mean cortical volume was positively associated with DMN (slope=1.66×10^−6^ ± 6.19 x10^−7^, t_66_=2.68, p=0.009, Figure 3b), subcortical (slope=1.62×10^−6^ ± 5.22 x10^−7^, t_66_=3.11, p=0.003, Figure 3b), medial prefrontal (slope=2.31×10^−6^ ± 8.17 x10^−7^, t_66_=2.83, p=0.006, Figure 3b), visual association (slope=2.18×10^−6^ ± 8.91 x10^−7^, t_66_=2.45, p=0.02, Figure 3b) and whole-brain (slope=1.60×10^−6^ ± 5.32 x10^−7^, t_66_=3.01, p=0.004, Figure 3b) connectome stability, indicating higher stability with larger brain volumes. No significant associations emerged between the network stabilities and longitudinal changes in cortical volume (all p>0.05). Finally, including hippocampus volume as a predictor in the models, revealed no association between mean or changes in hippocampus volume and connectome stability (all p>0.05).

Episodic memory performance was related to longitudinal connectome stability. The analyses revealed a significant negative association between connectome stability and mean episodic memory performance (i.e. the mean of PC1 across MRI_1_ and MRI_2_) for the subcortical network (slope=-0.05 ±0.02, t_68_=-2.92, p= 0.005, Figure 4a), indicating higher subcortical network stability with lower mean episodic memory performance. Similar analyses for the other networks revealed no significant associations after correcting for multiple comparisons. Furthermore, the analyses revealed a significant negative association between DMN network stability and change in episodic memory performance, indicating larger episodic memory decline between MRI_1_ and MRI_2_ in individuals with higher DMN stability (slope= −0.07 ±0.02, t_67_=-3.13, p=0.003, Figure 4b). In addition, there was a nominal significant association between changes in episodic memory performance and subcortical network stability (slope= −0.04 ±0.02, t_68_=-2.21, p=0.03). Finally, while mean cortical volume was associated with connectome stability, general linear models revealed no significant associations (all p>0.05) between cortical volume and mean or changes in PC1 while adjusting for age, sex, ICV and time between sessions.

**Figure 4.**
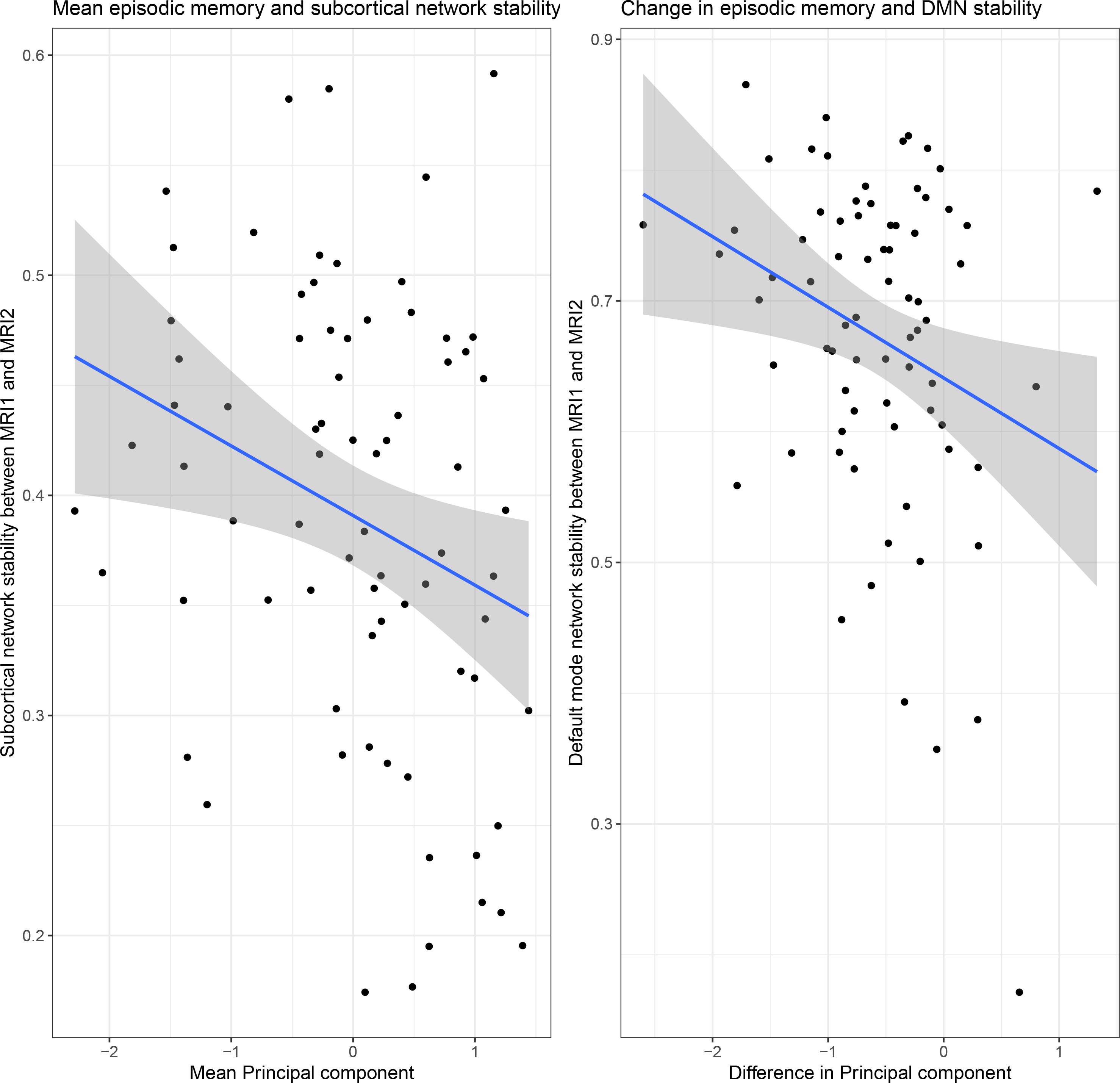
The association between episodic memory performance and subcortical or default mode network stability. (A) The association between mean (i.e. mean across MRI_1_ and MRI_2_) principal component 1 and subcortical network stability. (B) The association between individual changes in principal component 1 from MRI_1_ to MRI_2_ and default mode network stability.

## Discussion

In this study, we tested the long-term stability of the functional connectome in middle- and old age, and how brain network stability relates to structural indices of aging and memory performance. We demonstrated relatively high stability of the connectome over a 2-3 year time frame. While our analyses only revealed a subtle association with age, network stability was related to mean cortical volume across the two time points, suggesting that the structural integrity of the brain is associated with connectome stability. Supporting its relevance to cognitive aging, both cross-sectional and longitudinal measures of episodic memory were related to longitudinal stability of the DMN and the subcortical networks. The findings encourage the use of connectome stability for understanding individual differences related to brain aging and risk of neurodegenerative disease. Furthermore, the observation that individual variations in episodic memory decline relates to the stability of subcortical and DMN networks, provides new evidence for the importance of these networks in maintaining mnemonic processes in middle and old age.

Although previous studies have documented that the connectome individualizes during adolescence to form unique functional connectivity profiles (Kaufmann, Alnaes, Doan, et al., 2017), our knowledge regarding connectome stability in aging has been limited. The high stability obtained in the present study suggests that the connectome backbone may represent a robust trait-like marker also in middle and old age, despite the vast changes in brain structural and functional connectivity associated with increasing age. The finding is important as it opens a new avenue for studies of individual differences in cognitive aging as well as separating healthy aging from neurodegenerative disease. Despite large efforts in linking cognitive decline to brain changes in aging, the majority of previous studies have not been able to separate state- or task-based variability from static, subject-unique features. The notion that the connectome backbone generally remains stable across contexts and cognitive tasks (Finn et al., 2017; Kaufmann, Alnaes, Brandt, et al., 2017), and serves as a powerful predictor of cognitive abilities (Finn et al., 2015; Rosenberg et al., 2016), holds great promise for using connectome based approaches to map clinically useful changes of brain functional connections also in middle and old age.

Although the stability of the connectome backbone only showed a subtle association with age, the stability was related to individual differences in cortical volume. Aging is associated with structural degeneration of the cerebral cortex as indicated by widespread reductions in cortical thickness (Fjell, Westlye, et al., 2009) and grey matter volume (Good et al., 2001; Raz et al., 2005). However, individuals differ markedly in rate and degree of structural brain changes, and thus brain structure is relatively well preserved in some individuals into old age. Such individual differences is likely to be related to numerous factors, including individual variations in microvasculature (Dey, Stamenova, Turner, Black, & Levine, 2016) and white matter pathologies (Langen et al., 2017), processes which affects both local and distant brain connections and thus the connectome backbone. Of note, attempts to link cortical shrinkage to age related changes in cognition have been mixed (Salthouse, 2011) and was further supported by the lack of association between cortex volume and episodic memory performance in the present study. However, our finding that cortical volume relates to the connectome stability of medial prefrontal, DMN and subcortical networks suggests a mechanism by which cortical structural integrity could impact cognition. Thus, future studies may investigate if the association between cortical structure and age-related changes in cognition is mediated by the stability of the connectome, with consequences for our understanding of numerous neurodegenerative disorders.

The finding that the temporal stability of the subcortical network was inversely related to episodic memory performance may at first seem counterintuitive. However, increased subcortical network stability may not necessarily represent better brain maintenance. Increased network stability may come at the cost of decreased flexibility, including the task dependent engagement of cortico-subcortical networks in episodic memory tasks. In line with this notion, middle aged and older subjects experiencing increased striatal or hippocampal synchronization during rest, also had diminished cortical-subcortical connections and poorer memory performance in cross-sectional studies (Rieckmann et al., 2018; Salami et al., 2014). Previous studies have reported altered variability of neural networks with increasing age (Garrett, Kovacevic, McIntosh, & Grady, 2013; Guitart-Masip et al., 2016), which may depend on dopaminergic neurotransmission (Guitart-Masip et al., 2016). An optimal level of variability in neural activity and connectivity may increase detection of weak incoming signals (McDonnell & Ward, 2011) and also allow a greater range of responses to a greater range of stimuli (Deco, Jirsa, & McIntosh, 2011), all of which are likely to be important for episodic memory function. Moreover, the ability of neuronal networks to flexibly adapt in response to neurodegenerative changes, may be a prerequisite for maintaining cognitive function in older age. As such, increased subcortical within-network stability may reflect diminished ability to flexibly adapt in response to neurodegenerative changes locally and elsewhere in the brain, which translated into impaired episodic memory function.

In addition to the subcortical stability, subjects experiencing greater memory decline also had a more stable DMN during the 2-3 year time frame. This fits well with evidence that the DMN entails interacting subsystems that are implicated in episodic memory (Staffaroni et al., 2018), and that the longitudinal trajectory of DMN connectivity is associated with changes in episodic memory function in aging (Staffaroni et al., 2018). Moreover, increased DMN connectivity has been observed in mild cognitive impairment (Celone et al., 2006), preceding the profound reductions in whole-brain connectivity characteristic of Alzheimer disease. Although speculative, increased DMN stability may be a required permissive for the spread of pathological proteins, which eventually leads to aberrant network connectivity (de Haan, Mott, van Straaten, Scheltens, & Stam, 2012). In line with this heuristic, increased DMN synchrony over a lifetime is associated with total amyloid depositions in posterior DMN subsystems (Buckner et al., 2005). Moreover, the brains of healthy adults experiencing greater cognitive decline are more likely to harbor pathological proteins, including amyloid. Irrespective of this our results support the view that memory impairment in old age depends on simultaneous changes in multiple memory systems as connectome stability of different subnetworks (DMN, subcortical) was associated with episodic memory performance (Fjell et al., 2016). Accordingly, a whole-brain approach may provide a more holistic approach to memory in aging than the consideration of single networks or brain areas.

The vast majority of studies investigating resting-state functional connectivity in association to age-related memory changes have been cross-sectional. However, these studies do not allow determining whether the memory decline precedes the connectivity changes or the reverse. Among the few longitudinal exceptions, one study reported that the stability of the DMN was positively related to episodic memory maintenance in aging (Persson, Pudas, Nilsson, & Nyberg, 2014). Another study reported that better preservation of striatal-cortical connectivity over time yielded a more favorable memory outcome at follow-up testing (Fjell et al., 2016) possibly related to inhibition of subcortical intra-network connectivity at rest (Salami et al., 2014). While these studies investigated how individual changes in a common template of brain functional organization relates to episodic memory, they did not explore how age-related changes in the connectome backbone may affect memory function in old age. Accordingly, our finding that the stability of the subcortical and the DMN connectome relates to episodic memory in aging suggests that individual differences in the organization of subcortical and DMN networks influence how well mnemonic processes are maintained into old age.

The present study had some limitations. First, we note that the follow-up time of this study was relatively short, which may not be sufficient to detect reliable changes in functional connectivity in middle and older age. However, previous studies investigating longitudinal changes in functional brain connectivity in aging also used a follow-up time of approximately three years (Fjell et al., 2016). Moreover, grey matter atrophy (Storsve et al., 2014) and changes in structural connectivity (Sexton et al., 2014) could be reliably tracked over a three year time-interval, and possibly even shorter (Fjell, Walhovd, et al., 2009). Together, these studies suggest that a three-year period may be sufficient to detect age-related changes in functional connectivity. Moreover, we cannot rule out a practice effect in our subjects, which is a limitation in all longitudinal studies of neurocognitive aging. In the present study, an initial whole-brain approach was utilized, followed by predefined sub-network analyses. Although the subnetworks represent relatively coarse subdivisions of the brain, they do not exclude parts of the connectome, which is the case in all seed-based approaches. Thus, our findings should be followed-up using other approaches, including longitudinal seed-based studies investigating how alterations in the connectivity of specific subcortical nuclei correlate with age-related variations in memory.

In summary, our results suggest that the connectome backbone remains relatively stable over a 2-3 year time period in middle and old age. Individual differences in the stability of selected networks were associated with cortical volume and memory performance, supporting the neurocognitive relevance. Future large-scale longitudinal studies comprising genetics and rich cognitive and biological phenotyping with connectome-wide stability analyses could bring us closer to a mechanistic understanding of how age-related changes in neural events give rise to age-related cognitive decline, ranging from physiological changes to neurodegenerative disease.

## Aknowledgements

This work was supported by the Western Norway Health Authority (grant 911397 and 911687 to AJL, and grant 911593 to AL), and the Research Council of Norway (249795). The authors thank Dr. Jonn-Terje Geitung and Haraldplass Deaconess Hospital for access to MRI facility. The authors declare no conflict of interests.

